# Subcellular spatial proteomics of the close animal relative, *Capsaspora owczarzaki*

**DOI:** 10.64898/2026.10.13.758975

**Authors:** Dagmar Jirsová, Yu-Ping Poh, Katarzyna Dabrowska, Ritin Sharma, Patrick Pirrotte, Jeremy G. Wideman

## Abstract

*Capsaspora owczarzaki* is one of the closest relatives of animals and an increasingly tractable model for investigating the evolution of multicellularity. However, very little cell biological work has been done on the sister linages to animals. Subcellular spatial proteomics (SSP) is a technique that can localize thousands of proteins to subcellular compartments in a single experiment by combining cell fractionation with sensitive mass spectrometry. To provide broad insight into the localization of the *Capsaspora* proteome, we performed SSP based on a label-free version of LOPIT-DC (Localisation of Organelle Proteins by Isotope Tagging after Differential ultraCentrifugation). We detected 6,466 proteins, corresponding to 68.4% of the predicted *Capsaspora* proteome, and assigned high-confidence localizations to 3,106 of them across 13 subcellular compartments. This study serves as a proteomic dataset to resolve cellular compartmentalization in a close relative of animals, enabling downstream evolutionary, biological, and functional analyses. Investigators interested in using these data prior to publication are encouraged to contact us to avoid overlap and to facilitate coordinated and collaborative use of this resource.

## Introduction

The complexity of animal cells derives from the complexity of their genomes, life cycles, and many differentiated cell types. Unfortunately, unicellular models like yeast lack many of the parts and processes that occur in animal cells, human cell lines are immortalized and genetically abnormal, and primary cultures are short-lived and difficult to work with. A system that combines the tractability of yeast with the cell biology of animals is needed, and *Capsaspora owczarzaki* (*Capsaspora* henceforth) is a possible exemplar.

*Capsaspora* is an amoeboid protist, originally isolated as a symbiont of the freshwater snail *Biomphalaria glabrata* (Hertel et al., 2002), and one of the closest relatives of animals (Ruiz-Trillo et al., 2004; Torruella et al., 2015). It is a member of the Filasterea, which together with choanoflagellates and ichthyosporeans comprise the unicellular lineages of Holozoa (Shalchian-Tabrizi et al., 2008; Torruella et al., 2015). *Capsaspora* has three life stages: a proliferative filopodial amoeba that crawls on surfaces, an aggregative multicellular stage in which cells adhere to one another and secrete extracellular material, and a dormant cyst (Sebé-Pedrós et al., 2013). Aggregative multicellularity is regulated and can be induced by host-derived chemical cues (Ros-Rocher et al., 2023).

Work has shown that *Capsaspora* has many of the cellular features that are considered hallmarks of animal cell biology and development, making it an ideal candidate for investigating the evolution of multicellularity, animal cells, and cell types. Investigations into the genome and transcriptome of *Capsaspora* revealed hallmarks of animal multicellularity, including most of the integrin adhesion machinery (Sebé-Pedrós et al., 2010; Suga et al., 2013), components of GPCR signaling (de Mendoza et al., 2014), and transcription factors associated with animal development such as Brachyury, RUNX, and NF-κB (Sebé-Pedrós et al., 2011; de Mendoza et al., 2013; Suga et al., 2013). Its regulatory genome is remodeled between life stages (Sebé-Pedrós et al., 2016), and integrin-mediated adhesion is functional in *Capsaspora* cells (Parra-Acero et al., 2020). Though some features of Capsaspora appear fungal-like (Jirsová and Wideman, 2022; Ocaña-Pallarès et al., 2022), including the presence of the ER-mitochondria encounter structure (ERMES), which animals have lost (Wideman et al., 2013; Wideman and Muñoz-Gómez, 2016), its many animal-like features make it an excellent model system for investigating animal cell biology.

Although considerable effort has gone into understanding the development (via transcriptomics) and evolution (via comparative genomics) of *Capsaspora*, its cell biology has not been thoroughly explored, even though transfection and genome editing are now established (Parra-Acero et al., 2018; Phillips et al., 2022). Subcellular spatial proteomics (SSP) maps protein localization across the whole cell providing a kick-start to molecular cell biological analyses. Cells are gently lysed, organelles and other structures are separated into a series of fractions by centrifugation, and the abundance of each protein across the fractions is measured by quantitative mass spectrometry. Proteins that reside in the same compartment share a fractionation profile and can be assigned to a compartment by comparison to marker proteins with known location (Dunkley et al., 2004; Foster et al., 2006; Christoforou et al., 2016; Geladaki et al., 2019; Lundberg and Borner, 2019; Christopher et al., 2021; Breckels et al., 2024). First developed in plant and animal systems (Dunkley et al., 2006; Foster et al., 2006; Christoforou et al., 2016; Iuhak et al., 2017), SSP has since been applied across eukaryotic diversity, including many protists (Barylyuk et al., 2020; Guérin et al., 2023; Chisholm et al., 2026; Moloney et al., 2023; Hammond et al., 2026; Hammond et al., 2025; Jirsová et al., 2025; Chen et al., 2025; Zítek et al., 2022).

We use a label-free version of LOPIT-DC (Localisation of Organelle Proteins by Isotope Tagging after Differential ultraCentrifugation), in which fractions are generated by differential centrifugation (Geladaki et al., 2019). Here, we apply SSP to *Capsaspora* to generate the first subcellular proteome of a protist relative of animals. We detect 6,466 proteins, localize 3,106 of them to 13 subcellular compartments, and support these assignments with independent sequence-based predictions from TargetP (Almagro Armenteros et al., 2019), DeepTMHMM (Hallgren et al., 2022), and NLStradamus (Nguyen Ba et al., 2009). In sum, this work provides a new baseline for future cell biological investigations in this burgeoning model system.

## Results and Discussion

Capsaspora cells were grown axenically to ∼1 × 10^6^ cells/mL at room temperature (RT) of predominantly filopodial stage and harvested by centrifugation. Cells were gently lysed by four different approaches that preserve the integrity of intracellular organelles: nitrogen cavitation, dounce homogenization, and two different filter passaging methods. The resulting homogenate was separated into 11 fractions by differential centrifugation following the LOPIT-DC protocol with specific adjustments based on protein yield in individual fractions (Geladaki et al., 2019). The pellet from each spin, together with the supernatant remaining after the final spin, was collected for mass spectrometry, and the experiment was performed in quadruplicate. Protein recovery across the fractionation series was assessed by SDS-PAGE with Coomassie staining (Figure S1).

Label-free mass spectrometry analysis of 11 fractions, across four independent experiments, identified 6,466 proteins belonging to 6,211 protein groups. The identified proteome corresponds to 68.4% of the predicted *Capsaspora* proteome and recovers 88.2% of the eukaryota_odb10 BUSCO set (single-copy: 84.3%; duplicated: 3.9%). Protein abundances were normalized within each experimental replicate, and missing values were imputed by neighbour averaging followed by zeroing. Abundance profiles across the fractionation series were generated for each protein (Figure 1A) and visualized using t-distributed stochastic neighbor embedding (t-SNE) (van der Maaten and Hinton, 2008) to reveal protein clusters (Figure 1B).

**Figure 1.**
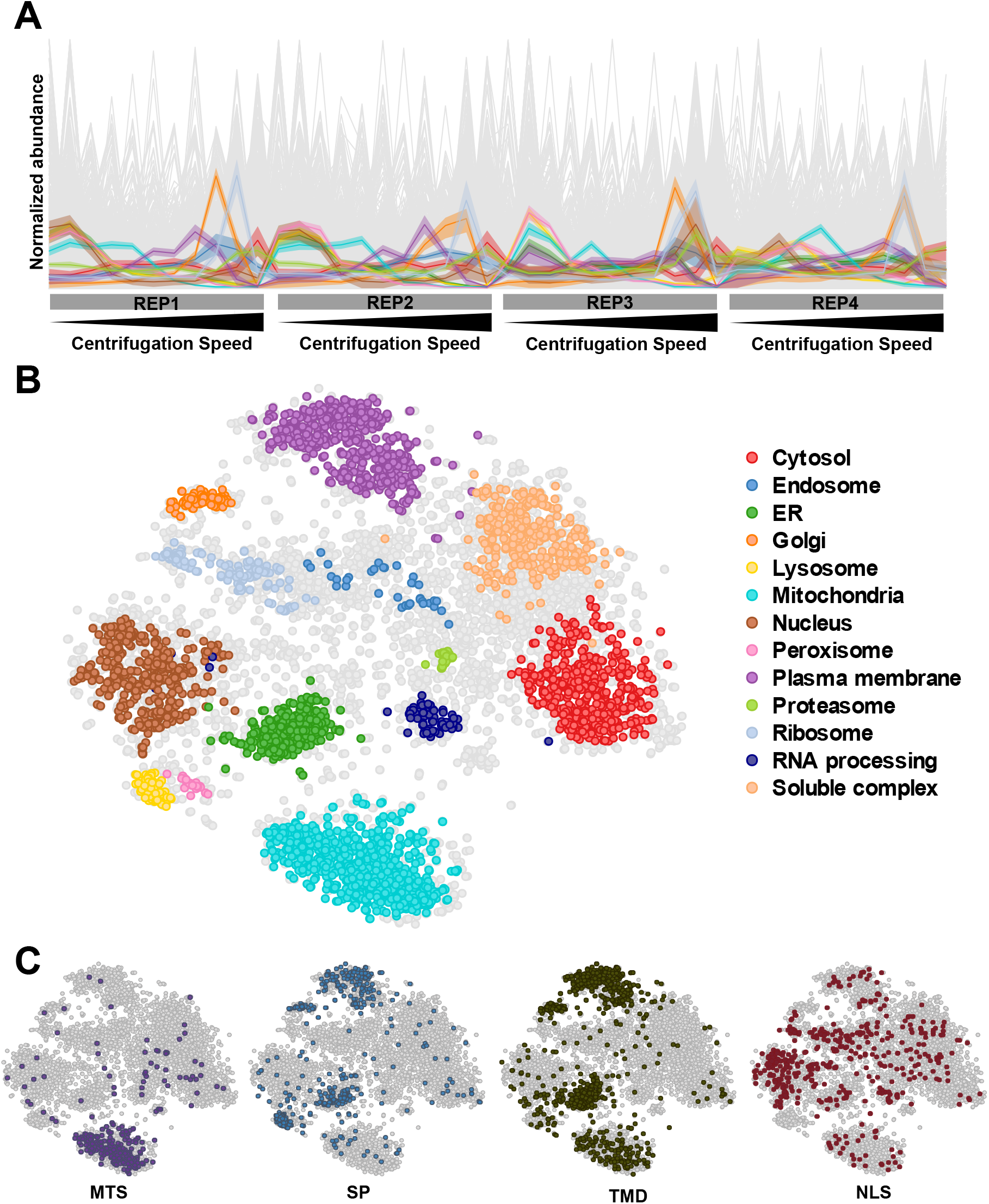
Subcellular proteome of *Capsaspora owczarzaki*. A. Normalized abundance profiles across the fractionation series for each of four independent experiments (REP1–REP4), with marker proteins coloured by compartment and all remaining proteins in grey. B. Protein localizations predicted by a supervised support vector machine classifier trained on 396 marker proteins across 13 compartments and filtered at a median probability cutoff, represented by t-SNE. C. Sequence-based targeting predictions mapped onto the same embedding, showing mitochondrial targeting signals (MTSs), signal peptides (SPs), transmembrane domains (TMDs) and nuclear localization signals (NLSs).

### Machine Learning-Based Assignment of Subcellular Localization

To evaluate the bias introduced by imputation, three datasets representing different levels of imputation were each classified by SVM using a manually curated marker set (Table S1). SVM assignments were consistent across the three datasets (Figure S2), and the dataset imputed by neighbour averaging followed by zeroing was therefore selected for downstream analysis in order to maximize protein coverage. Applying a median probability cutoff, the trained SVM assigned subcellular localizations to 3,106 *Capsaspora* proteins across 13 compartments (Figure 1B and Table S2), providing the first predicted localization for thousands of previously uncharacterized *Capsaspora* proteins.

To assess the reliability of the SVM assignments, we compared them against sequence-based prediction tools: TargetP (MTS and SP), DeepTMHMM (TMDs) and NLStradamus (NLSs). The predicted mitochondrial proteins formed a well-defined cluster in the t-SNE map, closely mirroring the distribution of proteins containing an MTS (Figure 1C). Furthermore, predicted endoplasmic reticulum (ER) and plasma membrane (PM) proteins mapped to the regions enriched for proteins with both transmembrane domains and signal peptides. In contrast, predicted lysosomal proteins fell within regions enriched for proteins carrying a signal peptide but lacking transmembrane helices. Overall, these sequence-level features strongly align with the predictions derived from our subcellular proteomics data.

For comparison, unsupervised k-means clustering with k = 13, matching the number of marker-defined compartments, yielded similar clustering patterns. However, the predicted cytosolic proteins were divided into three separate clusters, while the peroxisome and lysosome clusters merged into a single cluster (Figure S2).

## Conclusions

Here we report the first subcellular proteome of *Capsaspora owczarzaki*, and the first for any protist relative of animals. It provides an investigative resource for exploring its protein function, compartmental organization, and cellular architecture. Because many of the proteins that build animal cells predate animal multicellularity, knowing where those proteins localize in a unicellular holozoan is a prerequisite for interpreting what changed at the origin of animals. Investigators interested in using these data prior to publication are encouraged to contact the authors to avoid overlap and to facilitate coordinated and collaborative use of this resource.

### Data availability

The mass spectrometry proteomics data have been deposited to the ProteomeXchange Consortium via the PRIDE partner repository with the dataset identifier PXD085429 (Deutsch et al., 2020).

## Methods

### Culturing and Cell Harvesting

*Capsaspora owczarzaki* strain ATCC 30864 was cultured at room temperature (RT) in ATCC medium 1034 (modified PYNFH), containing 10 g/L Bacto peptone, 10 g/L yeast extract, 1 g/L yeast nucleic acid, 15 mg/L folic acid, 1 mg/L hemin, 2% (v/v) phosphate buffer (stock solution containing 18.1 g/L KH_2_PO_4_ and 25 g/L Na_2_HPO_4_), and 10% (v/v) heat-inactivated fetal bovine serum (FBS), adjusted to pH 6.5. Cultures were maintained by subculturing every two weeks and grown to a final cell density of approximately 1 × 10^6^ cells/mL in a total volume of 1.5 L. Cells were harvested after 3–4 days, during the filopodial stage, by centrifugation at 10,000 × g for 10 min at RT. The cell pellet was washed twice with 1× PBS, with centrifugation at 5,000 × g for 5 min after each wash. The washed pellet was resuspended in ice-cold detergent-free (DF) buffer (0.25 M sucrose, 10 mM HEPES, pH 7.4, and 2 mM EDTA) supplemented with Halt− Protease and Phosphatase Inhibitor Cocktail (Thermo Scientific) (PPIs), to a final volume of 8 mL, and subsequently subjected to cell lysis.

### Cell lysis and fractionation

Cells were gently disrupted using three different approaches, yielding four lysate preparations: (i) nitrogen cavitation; (ii) Dounce homogenization; and (iii) two membrane filtration variants. (i): The cell suspension was transferred into a cell disruption vessel (Parr 4639, Parr Instrument Co.) pre-chilled to 4 °C and connected to a nitrogen gas tank. The vessel was pressurized to 300 psi and equilibrated on ice for 15 min. After equilibration, the cell homogenate was collected through the outlet valve at approximately 3 drops/s, gently folding the foam into the liquid. (ii): Cells were lysed using a pre-chilled Dounce homogenizer equipped with a tight-fitting, B-type pestle. Briefly, 4 mL of cell suspension in DF buffer supplemented with protease and phosphatase inhibitors (PPIs) was transferred into the homogenizer and disrupted by 15 up-and-down strokes of the pestle. Lysis efficiency was evaluated by DAPI staining and microscopy. The lysate was cleared of unbroken cells, partially disrupted cells, and aggregates by centrifugation at 300 × g for 5 min at 4 °C, and the supernatant was kept on ice. The pellet was resuspended in 4 mL of DF buffer supplemented with PPIs, and the homogenization procedure was repeated to increase cell disruption efficiency. Both lysates were combined and subjected to differential centrifugation. (iii): Two different membrane filtration approaches were used for cell disruption. In the first approach, 8 mL of cell suspension was loaded into a syringe and passed through a stack of two overlaid 5.0 µm filter membranes (Millipore, SVLP02500) mounted in a Swinnex Filter Holder (Millipore, SX0002500). In the second approach, 8 mL of cell suspension was loaded into a syringe and passed through a single 5.0 µm membrane syringe filter (Millipore, SLSV0R25LS), followed by a second passage through a 1.2 µm membrane syringe filter (Minisart, 17593). Subcellular fractionation was performed according to a previously described protocol (Geladaki et al., 2019), with the following modifications. The original final centrifugation step at 120k × g/45 min yielded insufficient amount of protein for mass spectrometry analysis, while a substantial amount of protein remained in the supernatant. Therefore, the final centrifugation speed and duration were increased to 150k × g/60 min to improve protein recovery in the pellet. In the case of Dounce homogenization, centrifugation at both 120k × g and 150k × g yielded very little protein in the pellets, and therefore only the corresponding supernatant was used for downstream analysis. Differential centrifugation generated 11 fractions: ten successive pellets and the supernatant remaining after the final centrifugation step. One set of fractions was used to assess protein distribution by SDS-PAGE, while another was stored at −80 °C for subsequent mass spectrometry analysis. The experiment was performed in quadruplicate.

### SDS-PAGE and Coomassie Staining

Protein recovery and distribution across the fractionation series were assessed by SDS-PAGE. Pellets obtained from differential centrifugation were resuspended in 2× Laemmli buffer (4% SDS, 20% glycerol, 120 mM Tris-HCl pH 6.8) and heated at 95 °C until dissolved, and protein was quantified using the Pierce− BCA Protein Assay Kit (Thermo Scientific). 10 µg of protein per fraction was separated on Bolt Bis-Tris Plus Mini Protein Gels, 4–12% (1.0 mm, WedgeWell format, Invitrogen) alongside protein marker (Amersham ECL Rainbow Marker, Full Range). Gels were stained for 1 h in 0.25% Coomassie brilliant blue (Pierce Coomassie Brilliant Blue R-250, Thermo Scientific) in methanol/water/acetic acid (1:10), destained by diffusion in the same solution minus the dye, and stored in ddH2O to clear the background. Gels were imaged using an Azure 600 (Biosystems) (Figure S1).

### Proteomics of Capsaspora Organellar Fractions by LC-MS/MS

Pelleted organellar fractions were brought up in 500 µL lysis buffer (8M urea, 75mM NaCl, 1mM EDTA, 50mM Tris-HCl at pH 8.0, 1 x HALT Protease and Phosphatase Inhibitor Cocktail (Thermo Fisher Scientific)) and disrupted by sonication using four 15 s pulses at 50% amplitude, with 1 s intervals for a total of 1 min. Insoluble material was removed by centrifugation at 20,000 rcf for 10 min at 4°C. Total protein concentration in the clarified supernatant was determined using a BCA assay (Thermo). Aliquots of 50 µg protein from each sample were subjected to reduction with 5 mM dithiothreitol (DTT), followed by alkylation with 20 mM iodoacetamide (IAA). Samples were then digested overnight at 37°C using Trypsin Gold (MS grade, Promega) at an enzyme:substrate ratio of 1:25. Resulting peptides were purified by solid-phase extraction using 1cc 50 mg C18 SPE columns (Waters Seppak), dried under vacuum, and kept at -80°C until further use. Prior to analysis, dried peptides were resolubilized in 100 µL of 2% acetonitrile, 0.1% formic acid in water and their concentration was reassessed by BCA protein assay.

For spectral library construction, peptides from all samples were combined in equal proportions and subjected to high-pH reversed-phase HPLC fractionation, yielding 96 fractions. Separation was performed on a C18 column (XBridge, 3.5 µm, 4.6 × 100 mm, Waters) maintained at 30°C, with a flow rate of 0.5 mL/min across a 96 min gradient. The resulting 96 fractions were concatenated into 24 pooled fractions and taken to dryness under vacuum. Both individual samples and library fractions were resuspended in 2% acetonitrile, 0.1% formic acid in water supplemented with iRT peptides (Biognosys). LC-MS/MS acquisition was performed on a Vanquish Neo ultra high-performance liquid chromatography (UHPLC) system (Thermo Fisher Scientific) interfaced with an Orbitrap Ascend Tribrid mass spectrometer (Thermo Fisher Scientific) fitted with a FAIMS Pro device. For each injection, 5 µL of sample was loaded directly onto a C18 analytical column (EasySpray ES802A, 75 µm inner diameter × 25 cm, 2-µm particle size, 100 Å pore size) held at 45°C and eluted at 300 nL/min. Chromatographic separation employed a 120 min binary gradient using Solvent A (LC-MS grade water, 0.1% formic acid) and Solvent B (LC-MS grade acetonitrile, 0.1% formic acid). Spectral library fractions were measured in DDA mode with the following parameters: MS1 detection in the Orbitrap at 120K resolution, MS2 detection in the Orbitrap at 30K resolution, and HCD at an NCE of 25%. Individual samples were acquired in DIA mode with the following parameters: scan range of 380–980 m/z, 16 m/z isolation windows with 1 m/z overlap, Orbitrap detection at 15K resolution, HCD NCE of 25%, and FAIMS CV of -45. Sample-specific spectral library generation was carried out using the Pulsar search engine within Spectronaut (v19.0.240606.62635), with the following search parameters: the *Capsaspora owczarzaki* (strain ATCC 30864) reference proteome database downloaded from UniProt (https://www.uniprot.org/taxonomy/595528; Proteome ID UP000008743) was used as the target database, carbamidomethyl and oxidation were specified as peptide modifications, up to two missed cleavages were permitted, and both proteins and peptides were retained at a false discovery rate threshold below 1%.

DIA-MS data were then queried against the sample-specific spectral library in Spectronaut (v19.0.240606.62635) with default parameters, with the exception that cross normalization was disabled. Protein and peptide identifications were accepted at a 1% FDR threshold. Abundance values for proteins and peptides exported from Spectronaut were subsequently normalized by variance stabilizing normalization in R (v4.3.0). The resulting abundance profiles were imported into R (R Core Team, 2018) and analyzed using the Bioconductor packages MSnbase (v2.24.2; Gatto and Lilley, 2012) and pRoloc (v1.38.2; Gatto et al., 2014). The concatenated dataset from the four experiments yielded abundance profiles for 6,211 protein groups across 44 samples. The resulting table was imputed by neighbour averaging followed by zeroing. Principal component analysis and t-distributed stochastic neighbor embedding (t-SNE; van der Maaten and Hinton, 2008) were applied for dimensional reduction and data visualization.

### Supervised and Unsupervised Classification

A curated marker set of 396 proteins across 13 subcellular compartments (Table S1), each with previous experimental evidence of a single localization, was used to classify all 6,211 protein groups. To classify all the proteins into the subcellular compartments, we first used supervised classification by support vector machine (SVM) model with the functions, svmOptimization and svmClassification, implemented in pRoloc (Gatto et al., 2014). For all clusters, the global median was used to set the cutoff score; proteins with an SVM score above the cutoff were considered predicted to their corresponding compartment, and those below were reset to “unknown”. The t-SNE map of the SVM-predicted proteins was plotted using the Shiny application implemented in pRolocVis (v1.38.2).

In parallel, unsupervised k-means clustering was performed using the MLearn function from MLInterfaces (v1.78.0) in R to generate 13 clusters, within which protein distribution profiles are more similar to one another than to those of other clusters.

### Targeting sequence identification

TargetP 2.0 (Almagro Armenteros et al., 2019) was used to predict SPs and N-terminal MTSs. The organism group was set to “Plant” to prevent mammalian-specific parameter bias. DeepTMHMM v1.0.24 (Hallgren et al., 2022) was applied to predict transmembrane topology. Finally, nuclear localization signals (NLS) were identified using NLStradamus (Nguyen Ba et al., 2009) with the two-state static HMM and a posterior probability cutoff of 0.95.

## Acknowledgements

J.G.W. was supported by a grant from the National Science Foundation (DBI-2119963) and a grant from the Gordon and Betty Moore Foundation (GBMF10600). D.J. was supported by VEDA FELLOWSHIPS within the Operational program Jan Amos Komensky (OP JAK), Project No. CZ.02.01.01/00/22_010/0008117. This research includes work conducted in the Integrated Mass Spectrometry Shared Resource supported by the National Cancer Institute of the NIH under grant number P30CA033572.

## Supporting Information

**Table S1. Marker proteins used in this study**.

**Table S2. SVM scores for all proteins detected in this study**.

**Figure S1.**
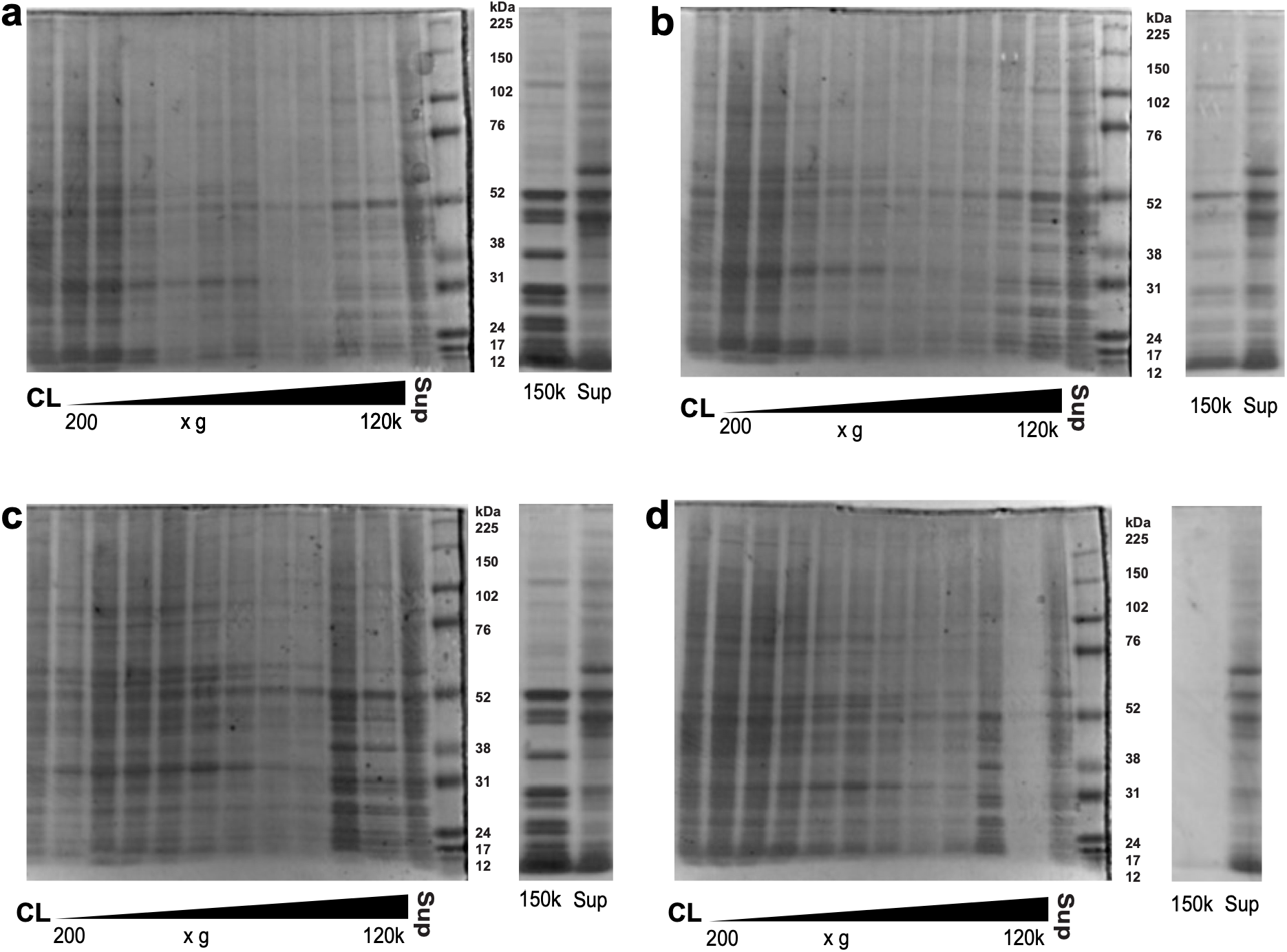
Coomassie-stained SDS-PAGE of subcellular fractions of *Capsaspora owczarzaki*. Fractions are arranged in order of increasing centrifugation speed, with cell lysate and the post-ultracentrifugation supernatant in the first and last lanes. We used four lysate preparations: (A) nitrogen cavitation (300 psi/15 min);(B) passing through a stack of two 5.0 µm filters; (C) passage through a 5.0 µm membrane filter followed by a 1.2 µm membrane syringe filter; and (D) Dounce homogenization. Differences in band pattern across the series indicate that protein was partitioned between fractions rather than carried through the series uniformly.

**Figure S2.**
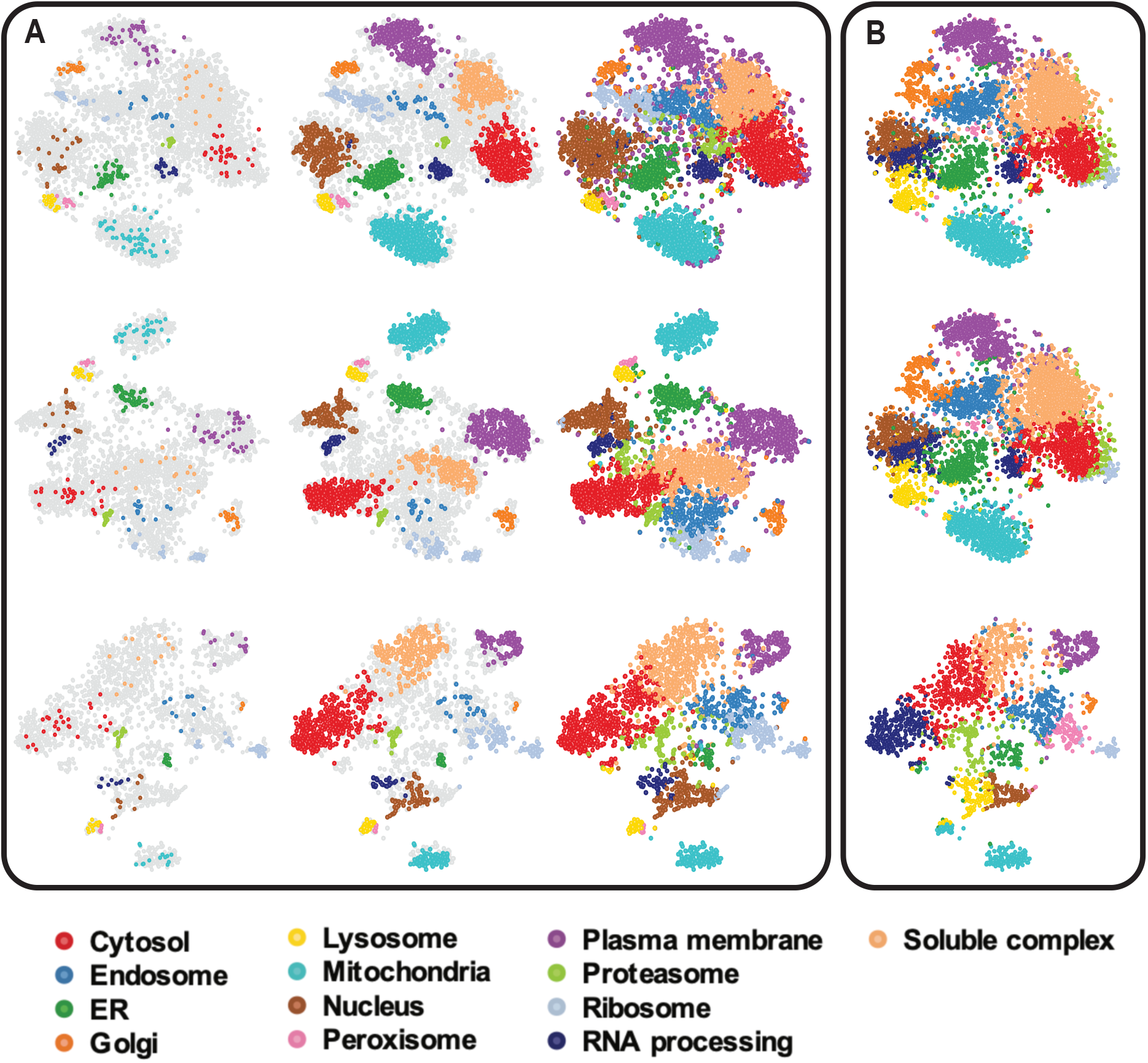
Subcellular atlases of *Capsaspora owczarzaki* under different imputation strategies and k-means clustering. A. SVM classification of datasets with neighbour averaging followed by zeroing (top row), neighbour averaging (middle row), and no imputation (bottom row). For each, the marker set, the assignments retained at the median SVM cutoff, and all assignments with no cutoff are shown from left to right. B. Unsupervised k-means clustering with k = 13, for the same three imputation strategies. k-means clusters have been given the colour of the marker-defined compartment to which they most closely correspond.

